# A Local Hebbian Rule Based Neural Network Model of Invariant Object Representation and Classification

**DOI:** 10.1101/2022.10.14.511519

**Authors:** Rishabh Raj, C. Ron Yu

## Abstract

Our recognition of an object is consistent across conditions, unaffected by motion, perspective, rotation, and corruption. This robustness is thought to be enabled by invariant object representations, but how the brain achieves it remains unknown^1^. In artificial neural networks, learning to represent objects is simulated as an optimization process^2^. The system reduces discrepancies between actual and desired outputs by updating specific connections through mechanisms such as error backpropagation^3^. These operations are biologically implausible primarily because they require individual connections at all levels to be sensitive to errors found at the late stages of the network^4,5^. On the other hand, learning in the nervous system occurs locally, and synaptic changes depend only on pre- and post-synaptic activities^6,7^. It is unclear how local updates translate into coordinated changes across large populations of neurons and lead to sophisticated cognitive functions. Here we demonstrate that it is possible to achieve robust and invariant object representations in naturally observed network architectures using only biologically realistic local learning rules. Adopting operations fundamentally different from current ANN models, unsupervised recurrent networks can learn to represent and categorize objects through sensory experiences without propagating or detecting errors. This white box, fully interpretable networks can extract clean images from their corrupted forms and produce representations prospectively robust against unfamiliar perturbations. Continuous learning does not cause catastrophic forgetting commonly observed in ANNs. Without explicit instructions, the networks can classify objects and represent the identity of 3D objects regardless of perspective, size, or position. These findings have substantial implications for understanding how biological brains achieve invariant object representation and for developing biologically realistic intelligent networks that are efficient and robust.

## INTRODUCTION

Sensory stimuli evoke high-dimensional neuronal activities that reflect not only the identities of different objects but also context, the brain’s internal state, and other sensorimotor activities^8^. The high-dimensional responses can be mapped to object-specific low-dimensional manifolds that remain unperturbed by neuronal and environmental variability^9,10^. It is not known how the brain generates an invariant representation of the same object not only in the face of signal corruption but also against variations in size, location, and perspective. Early computer vision models relied on convolutions and serial integration of features to produce size and location invariant object representations^11–16^. These models successfully explained specific aspects of visual processing, and later models using deep artificial neural networks (ANNs) mimic cognitive functions and show remarkable performances^17,18^. The successes rely on a learning process that minimizes discrepancy (error) between the desired output and the output produced by the network. During training, the network must “know” pre-determined sets of inputs and their corresponding outcomes^19^. Any detected mismatch is propagated throughout the network to update the connections to minimize the error^20,21^. The goal-directed updates and supervised training make the network exceptionally accurate in performing specific tasks. However, this striking accuracy comes at various costs. For example, the network cannot learn indefinitely. Upon completion of training, the updated connection weights are “frozen” and do not change further; exposition to new tasks can lead to catastrophic forgetting^22–24^. Training on specific examples does not generalize well beyond its training data^25–28^ and also makes the networks vulnerable to adversarial attacks^29–32^. To improve performance and robustness, numerous layers and large amounts of training data are required.

Although it has been suggested that the ANN models recapitulate computations taking place in the brain^33^, the nervous system operates differently. Biological brains do not know specific inputs a *priori*. They learn without instructions or labels, and there is no natural mechanism to back-propagate errors^34^. An essential aspect of biological systems is local computations^35–38^. Modifications in the synaptic strengths are instructed only by the activities of pre-and post-synaptic neurons, indifferent to changes in other brain parts^39–45^. The organic system is also constantly updated through experience, and unlike ANNs, they are remarkably robust against the kind of adversarial attacks to ANNs^46–48^. To be biologically realistic, artificial network models must use local learning rules to achieve global success in representing and classifying objects.

Since synaptic changes are oblivious to other parts of the neural network, global changes must arise from how neurons are connected. Here we present two network configurations that capture dependencies among features belonging to the same object and achieve consistent object representation and classification using solely local learning rules^49^. A discriminative network layout produces unique representations of objects robust against signal corruption and occlusion. An added classification module enables highly accurate categorization in a completely unsupervised manner. The two-module network can also represent the identity of 3D objects independent of viewing perspectives, size, or position. Notably, the network is not ostensibly designed to accomplish these goals. Nor does it perform error detection or propagation to achieve gradient descent. These examples demonstrate how specific architectures combined with local rules can enable networks to function like a biological brain and to be highly efficient and robust in processing information.

## RESULTS

### A dependency capturing network for object discrimination and representation

We sought a network architecture that allows individual neurons to capture maximum information about distinct objects and can consistently represent an image^49^, meaning that the network responds with the same activity pattern to an image and its various corrupted forms. We constructed a two-layer network. The first layer (input layer) detects and transmits the input patterns and projects to the second recurrent layer, the discrimination layer, in an all-to-all manner. Recurrent connections within the discrimination layer are also all-to-all and are inhibitory (Fig. 1a). Although the recurrency is reminiscent of the Hopfield networks^50,51^, this configuration generates sparse and distinctive activity patterns for each input^52^.

**Fig. 1.**
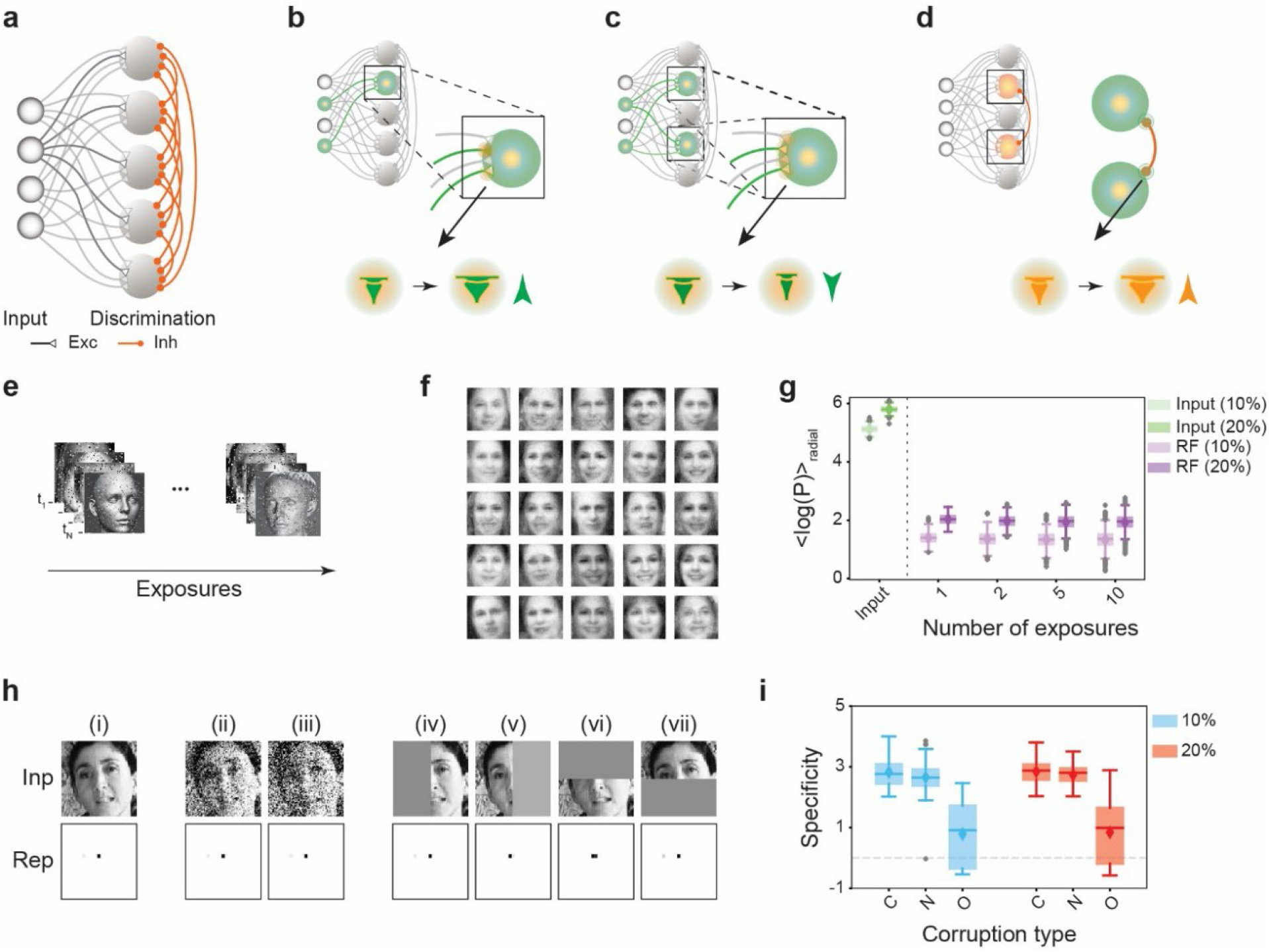
A network configuration to capture informative structures. **a.** A schematic representation of the network. A recurrent network of neurons constitutes the discrimination layer and receives excitatory connection from input neurons. The recurrent and the input connections are all-to-all. Recurrence is inhibitory (red connections). **b-d**. Local learning rules: the connection between an input neuron and a discrimination neuron strengthens when they are both active (**b**; green, synapses highlighted in yellow); when two neurons in the discrimination layer are co-active, all input connections to these neurons are weakened (**c**); reciprocal inhibitory weights increase when two discrimination layer neurons have the same activity (**d**). The updated synapses are highlighted in the inset. **e.** The network learns facial images corrupted by random noise from sequential exposure. The same image appearing in different baches was corrupted by different noise patterns. **f.** Examples of input structures captured by the network when exposed to faces with 10% noise. Note that the learned structures or the receptive fields (RFs) resemble complete faces but are not specific to any input image. **g.** The average power in the highest spatial frequencies in the noise-corrupted images and in the RFs. Learned RFs have much lower noise level. **h.** Representation (Rep) in the discrimination module of a previously unseen face (Inp i) and its corrupted forms by Gaussian noise (Inp ii-iii) or occlusions (Inp, iv-vii). The representations (Reps) are consistent with each other. **i.** Representation specificity of 100 distinct clean faces (C) compared to the corresponding values for their noise corrupted (N) and occluded (O) variations. The specificities of noisy examples match the clean images in both learning conditions. Occluded forms show lower specificity but are significantly higher than non-specific representations (dotted line). Images in **e** and **f** are from ref ^49^.

A distinguishing feature of our network is that the initial connectivity between the input and the discrimination layers takes the variance structure of the input dataset into account and ensures that two neurons are less likely to fire together for any input (see methods). Moreover, the learning process does not utilize any label, nor with any pre-determined outcome. It is entirely unsupervised, as the representations evolve with exposures to individual images. Thus, the recurrent weights do not reflect the correlation structure between pre-determined representation patterns. Importantly, the learning rules are all local and modeled as the following.

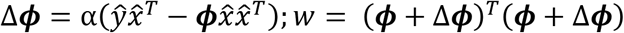

where *ŷ* is an input vector, 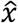 is its representation in the discrimination layer, ***ϕ*** is the connectivity between the input and the discrimination layer, α is the learning rate, and *w* is the recurrent inhibition weight matrix. The updates enable the network to learn comprehensive input structures without resorting to using reconstruction error or credit assignment^20,21,53^ (Methods). This formulation can be interpreted as separate rules at three synapses. Concurrent with the linear sum of activities to drive responses, the network adjusts connection strengths in an activity-dependent manner. The first term of the learning rule is a small increment of the connection strengthens when both input and discrimination layer neurons are active (Fig. 1b). This update allows the association between a feature (in the input) with the representation unit to capture the information^54^. The second term indicates that when two neurons in the recurrent layer are co-active (and mutually inhibited), the strengths of all synapses from the input to these neurons are reduced (Fig. 1c). The inhibitory weights in the recurrent layer are such that neurons responding to similar inputs have strong mutual inhibition (Fig. 1d). These updates essentially local Hebbian or anti-Hebbian rules, where synaptic updates are solely determined by neural activity. This configuration, i.e., the initial biased connectivity and local learning rules, distinguish our model from others, which incorporate random initial connections from the input that do not update (e.g., the convolutional input strengths in the ANN models). Note that all activities in the neurons and the synapses are non-negative to reflect the biological constraints.

We tested the performance of this network. A vital feature of the biological system is that it can learn from noisy examples. To simulate this process, we exposed the network to a series of images corrupted by noise during training. For example, we added random salt-and-pepper noise to facial images and fed them sequentially to the network (Fig. 1e). Note that we did not provide any expected responses for comparison purposes and there was no specific cost function associated with the learning process. As such, there was no error detection nor propagation. Following a series of exposures, we examined the structures learned by the network, or the receptive fields of the representation neurons. The structures resembled faces but were not specific to any input face (Fig.1f, Extended Data Fig. 1). Interestingly, the receptive fields appeared much less noisy than the input faces. We measured noise content in the input images and the receptive fields as the average power in the highest spatial frequencies. A higher mean power indicated higher noise content. We found that noise in the receptive fields was considerably lower than the input at all levels of training (Fig. 1g). This demonstrated that the network could inherently denoise the inputs and extract cleaner structures.

We tested the ability of the network to represent face images not in the training set, including unseen pictures corrupted by Gaussian noises or with occlusions. In these cases, the network generated sparse and consistent representations of the new faces. Representation of corrupted inputs were nearly identical to that of the clean images (Fig. 1h). Even images with large occlusion represented consistently (Fig. 1h iv-vii). By characterizing the representation performance, we found specificity remained high for corruptions with all noise levels and occlusions (Fig. 1i). This result highlights the network’s ability to learn from pure experience and generate consistent representations. Importantly, it achieves prospective robustness, defined as consistently representing input patterns it has never experienced^49^.

### Rapid receptive fields development and continuous learning

To understand synaptic weight changes during learning, we analyzed the receptive field properties of the recurrent layer neurons. The network trained on specific set of images rapidly learns the receptive fields that conform to the images (Figs.1f, 2a; Extended Data Fig. 1). For example, in a network trained using symbols from world languages, similarity between the receptive fields and the symbols increased rapidly as the network repeatedly encountered the same characters (Figs. 2a and 2b). The specificity of symbols’ representations increased even faster, reaching the plateau with less than 10 exposures (Fig. 2c). Thus, the network effectively captures structural features that are maximally informative about the input.

**Fig. 2.**
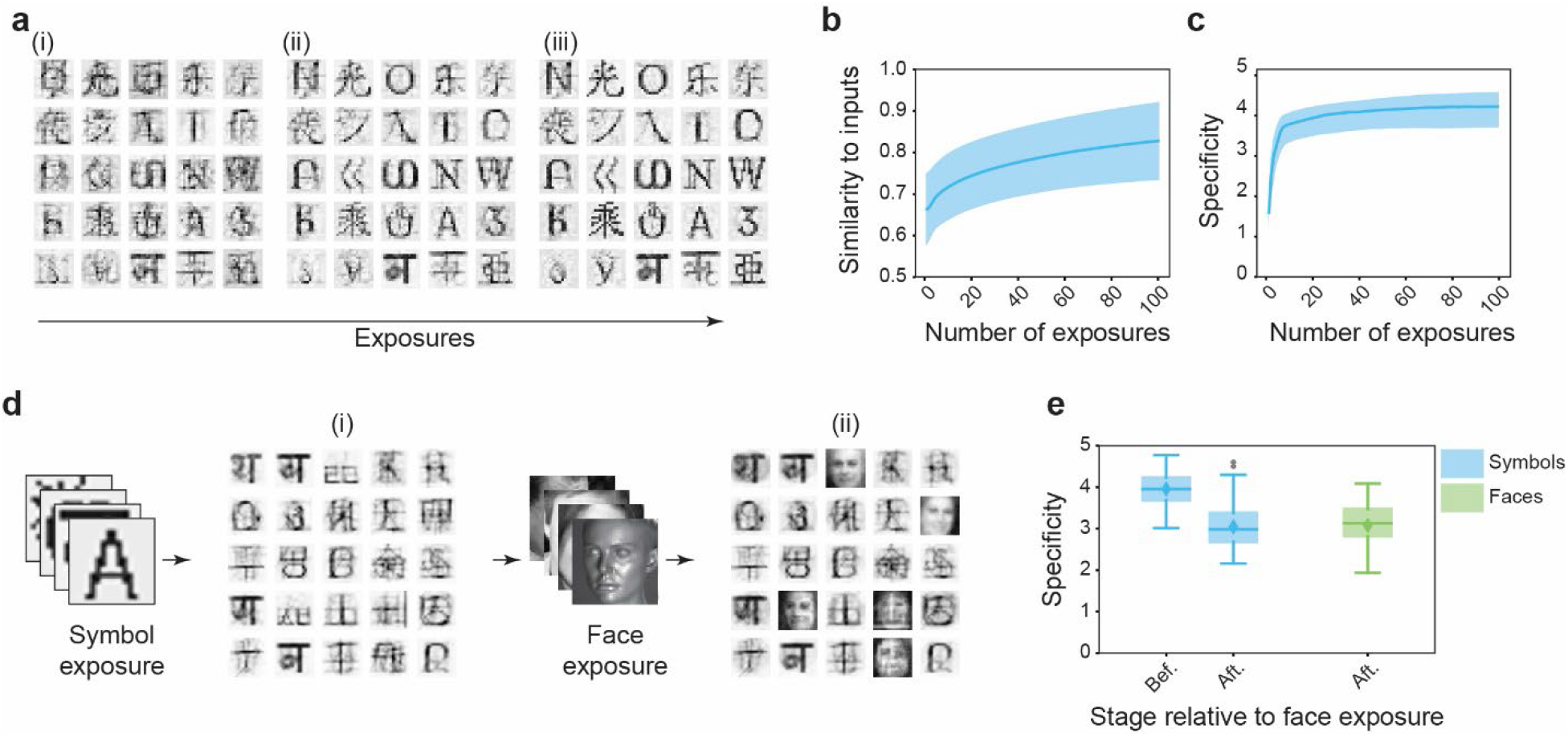
Rapid development of receptive fields and continuous learning in the network. **a.** Examples of RFs developed in the network from experiences of symbols from world languages. The RFs become clear gradually after 5 (i), 50 (ii), and 100 (iii) exposures to the symbols. **b.** Similarity of RFs to input images as a function of repeated exposures. **c.** Specificity of symbol representations increases as a function of exposures. **d.** Receptive field development following exposure to two different image sets sequentially. (i) The RFs after symbol exposure. (ii). RFs following subsequent exposure to faces. RFs resembling symbols remain whereas RFs resembling faces develop. **e.** The specificities of symbol representations before and after exposure to faces, which are comparable to each other. The representation specificity of faces is also high. Symbols and images are from ref ^49^.

Next, we tested whether the network could learn to represent novel input types without compromising its previous discrimination abilities. We first trained the network to represent a fixed set of symbols, followed by learning faces. Learning faces after the characters changed the receptive field properties of a subset of neurons (Fig. 2d). Remarkably, the specificity of symbol representations before and after learning the faces remained comparably high. The network also maintained high specificity of face representations (Fig. 2e). The representations remained consistent when the sequence was reversed, learning symbols followed learning faces (Extended Data Fig. 2). Thus, by maintaining its ability to represent previously learned objects while accommodating new ones, the network avoided the catastrophic forgetting problem encountered by the many ANN models^55,56^.

### Redundancy capturing architecture for classification

The discrimination module maximizes differences between objects and represents them distinctively. For classification, the network must capture shared features that identify an object in different perspectives, or a class. We reasoned that the distinguishing features of the same type of objects could be linked together using mutual excitation and discerned from similar features of other categories using inhibition. In the vertebrate brain, recurrent excitation and broad inhibition are prevalent in the upper layers of sensory cortices. We devised a recurrent network, the classification layer, to simulate these circuit motifs and perform computations for classification (Fig. 3a). Neurons in this module receive direct excitatory input from the discrimination module in a column-like, one-to-one manner. In parallel, they receive feedforward inhibitions that mirror the excitatory input. The neurons in the classification layer also have recurrent excitatory connections and receive global inhibition imposed on all neurons. The learning rule is that the synapses strengthen between two excitatory neurons (discrimination to classification and between classification neurons) when both are active (Fig 3b-c). There is no weight change to synapse to and from inhibitory neurons.

**Fig 3.**
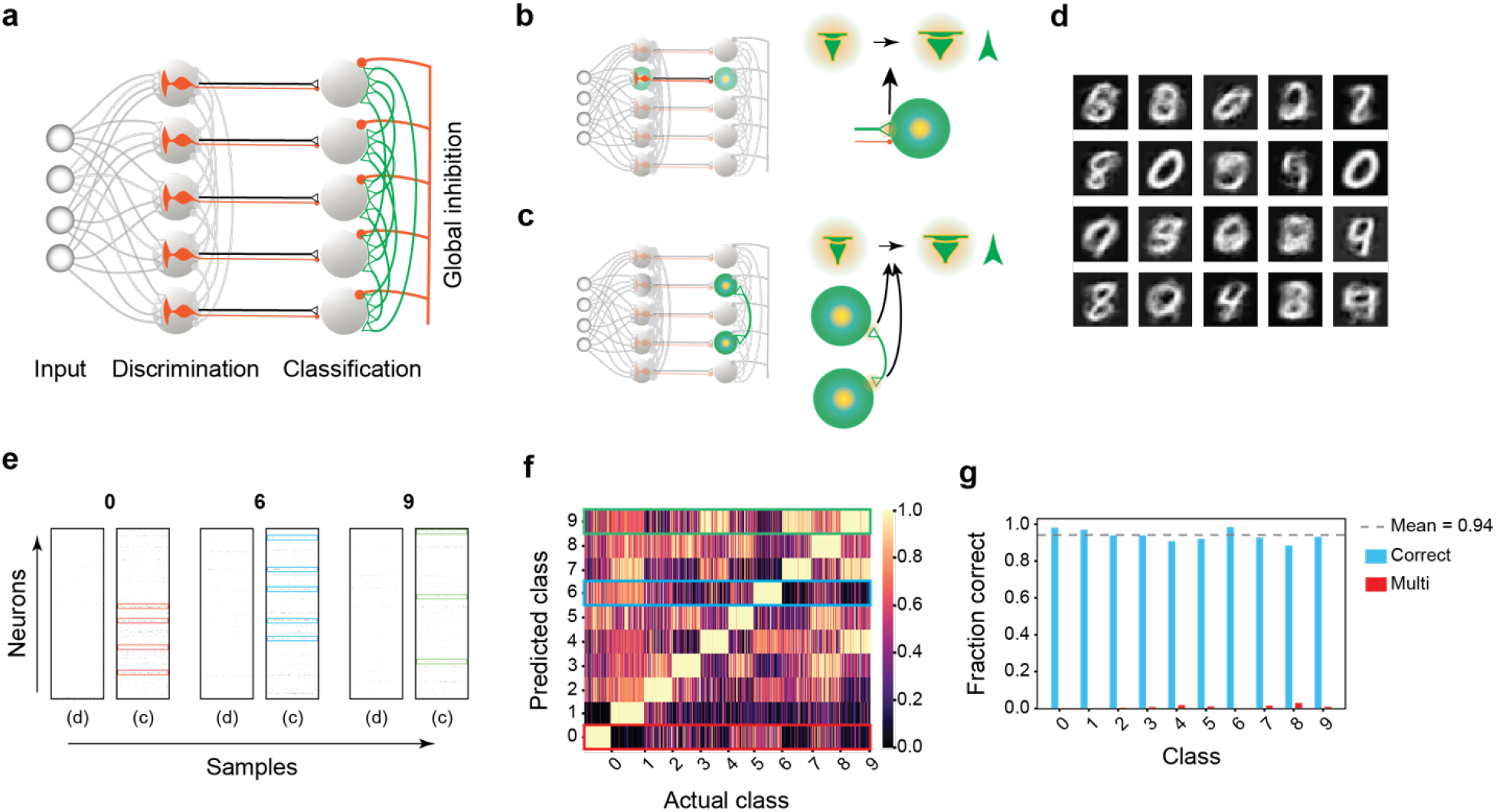
A network configuration for classification. **a.** Illustration of network configuration of the classification module. Each neuron receives one-to-one excitatory input from the discrimination layer and a one-to-one inhibitory connection from input neurons. The inhibitory neuron (red) receives the same input as the excitatory one but do not update synaptic weight. Neurons in the classification module receive reciprocal excitation (green) and global inhibition (red). **b and c.** Learning rules in the classification module. When a discrimination and a classification neuron are both active (b), or two neurons in the classification module are co-active (c), synaptic weights are augmented. Inhibitory connections to classification neurons, either from input neurons or from global inhibition, do not change. **d.** The discrimination layer extracts complete digit structures from images of handwritten digits in the MNIST dataset. **e.** The responses of neurons (ticks) in the discrimination (d) and classification (c) layers to examples of classes 0, 6, and 9. Responses to the digits in the discrimination module are sparse. Cells in the classification module consistently respond to all examples of the same digit type are highlighted in red (class 0), blue (class 6), and green (class 9). **f.** Heatmap of pooled responses (normalized to maximal activity) from neurons in the classification layer. The highest activities are for respective classes. Classes 0, 6, and 9 are highlighted in red, blue, and green boxes, respectively. **g.** Classification accuracy computed using pooled activity is high for all digit types (blue bars, mean ~94%). Some examples are classified into more than one class (red bars), but their proportion is negligible.

This architectural configuration permits capturing class-specific features from objects. First, neurons in the classification layer receive excitatory input from the discrimination module and feedforward inhibition relayed from the input layer. This combination passes the difference between the updated excitatory output and non-updated inhibitory output to informs the classification layer about the features learned in the discrimination layer. Then, the lateral excitatory connection between the classification neurons links the correlated features that provide the class information. Finally, global inhibition ensures that only neurons receiving sufficient excitatory input can be active to reduce spurious and runaway activities. The result is that neurons with reciprocal excitation display attractor-like activities for class-specific features.

We tested the network’s ability to classify objects using the MNIST handwritten digit dataset. Training with only 25% of unlabeled samples resulted in receptive fields resembling the digits in the discrimination module (Fig. 3d). Population activities in the classification module exhibited high concordance for the same digit type but maintained distinction among different classes (Fig. 3e). We identified the consistently active neurons in each category and pooled their activities. Interestingly, the pooled activities were highest for the respective groups (Fig. 3f). Therefore, we utilized them to evaluate classification accuracy. When tested using 5000 new samples, the network correctly identified 94% of the digit types (Fig. 3g). The most sophisticated network models currently achieve 85-99% accuracy, but they all need supervision in some form. For example, the self-supervised networks require digit labels in the initial training^57^.

### Extracting object identity from view perspectives

Arguably, the structure of biological brains allows them to accommodate diverse functions, whereas the ANNs perform only specific tasks. We have shown that the network is robust in recognizing and categorizing individual symbols, faces, and handwritten digits without explicitly designing it for the tasks. Specifically, in its discrimination module, the network can identify features that uniquely identify an object and, in the classification module, link those features to form class-specific neuronal ensembles. However, for a 3-dimensional object, views vary in size, position, and perspective. Relating them to extract the object’s identity is particularly challenging. Various ANN models require highly sophisticated algorithms with deep convolution layers and considerable supervision to achieve good performance^58–60^. We reasoned that different views of the same object form an image class that has shared features, which our network may capture without ostensibly being designed to do so. Therefore, we tested whether the network could learn to consistently represent 3D object images varying in size, position, and perspective.

To assess representational consistency against variations in size and position, we animated 50 different object models that vary in size or move across a field of view. We let the network experience several short clips as contiguous movie frames to simulate a real-life situation. These random clips could be partially overlapped but covered less than 33% of the entire animation sequence in total (Fig. 4a). Without definite instructions, the network learned specific views and superpositions of the objects (Fig. 4b-c). After training, we tested how well the network represented the object using the entire animation sequence, which contained sizes or positions that it had never seen (> 67% of all views). While the representations of different frames were distinct in the discrimination module, we observed cells persistently active over large animation portions in the classification module (Fig. 4d). Such consistent responses were observed for all objects. Importantly, active neuronal ensembles were specific for individual objects even when there were high similarities between some of them (Fig. 4d). As a result, in the representation domain, the overall similarity between the same object’s views was significantly higher than the similarity between images of distinct objects (Fig. 4e).

**Fig. 4.**
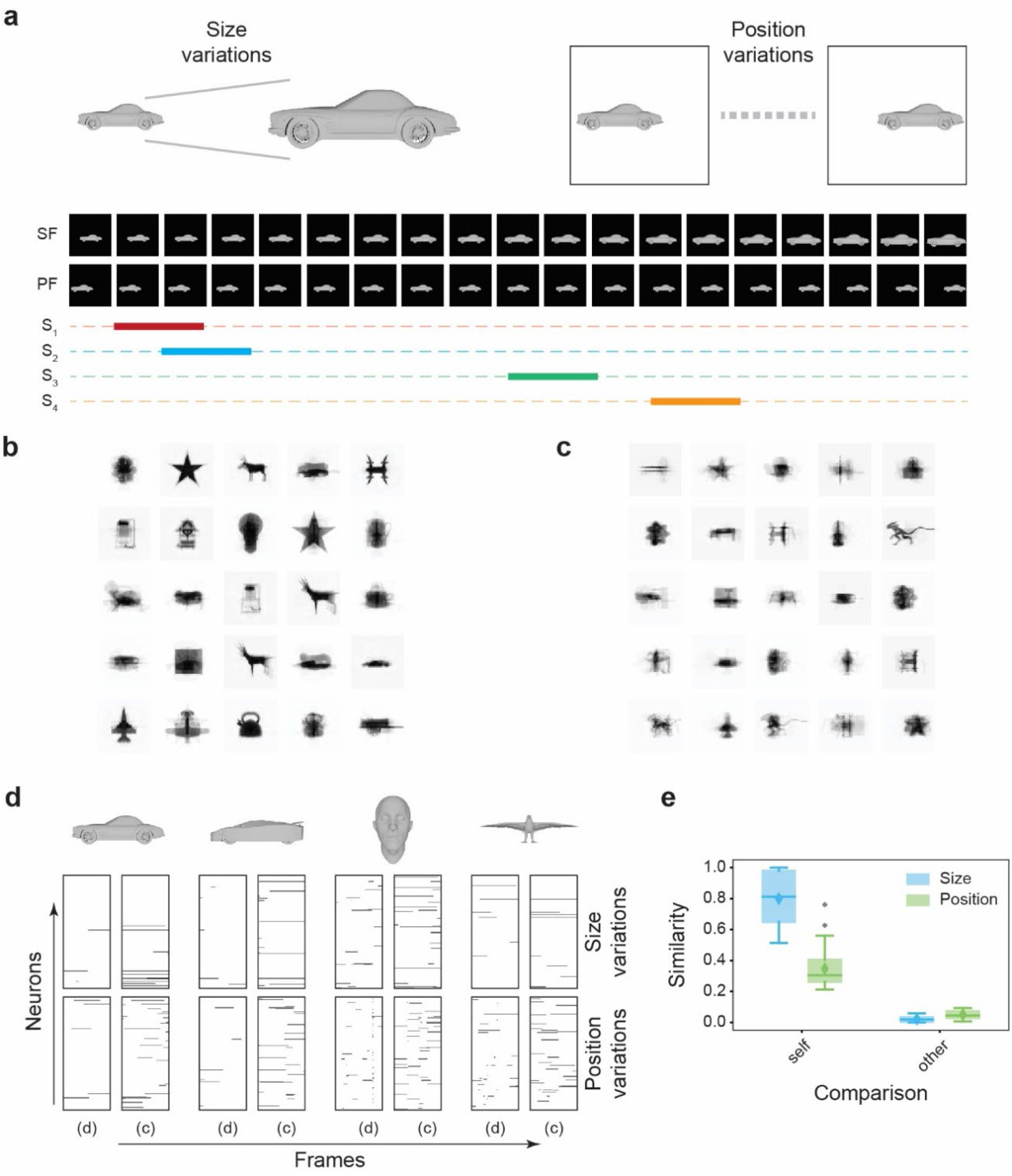
Size and position invariant representations of objects. **a.** Animating variations in size and position. The animations were rendered as movie frames depicting the size variations (SF) and position variations (PF). Short sequences of these frames (shown as solid, colored bars in S1-S4) covering not more than 33% of the entire sequence in total were randomly selected and fed into the network. **b-c.** In the discrimination module, the network captures complete object shapes varying in sizes (**b**) and positions (**c**). **d.** The responses of the discrimination (d) and classification (c) layers to all animation frames of different objects. The discrimination responses are discrete, whereas in the classification layers, continuous responses are observed for large portions of size and position-varying animations. Note that distinct neuronal ensembles show consistent responses in the classification layer for similar objects. **e.** The average similarities between representations of frames belonging to the same object (self) are considerably higher than the representation similarities between frames of distinct objects (other).

In these simulations, an object’s orientation remained fixed when it varied in size or changed position. Disparate perspectives arise when the object rotates along an axis. Producing representation invariant to 3D rotations is a challenging task^61^. To test whether our network could connect different 3D perspectives to produce identity representations, we generated animation of 3D rotation sequences and trained the network on short clips of rotation along the vertical axis (Fig. 5a). We observed view-specific receptive fields and responses in the discrimination module (Fig. 5b). In the classification module, cells showed consistent responses to the same object regardless of the presentation angle. Population responses to highly similar objects were distinct (Fig. 5c). Using a measure of representation consistency (Extended Data Fig. 3) we found smooth representation of the objects even for highly irregular shaped models (Fig. 5d). For the four 4-legged animals, fluctuations in representations of occurred at similar viewpoints, reflecting their common features. Overall, the similarity between the different perspectives of the same object was high but low between different objects (Fig. 5e). These results, in combination with the previous, highlight the network’s ability to generate invariant identity representations. Note that the network only experiences <33% of all possible angles. This, the capacity for invariant representation is prospective, meaning the network does not need to encounter all possible variations to represent them consistently^49^.

**Fig. 5.**
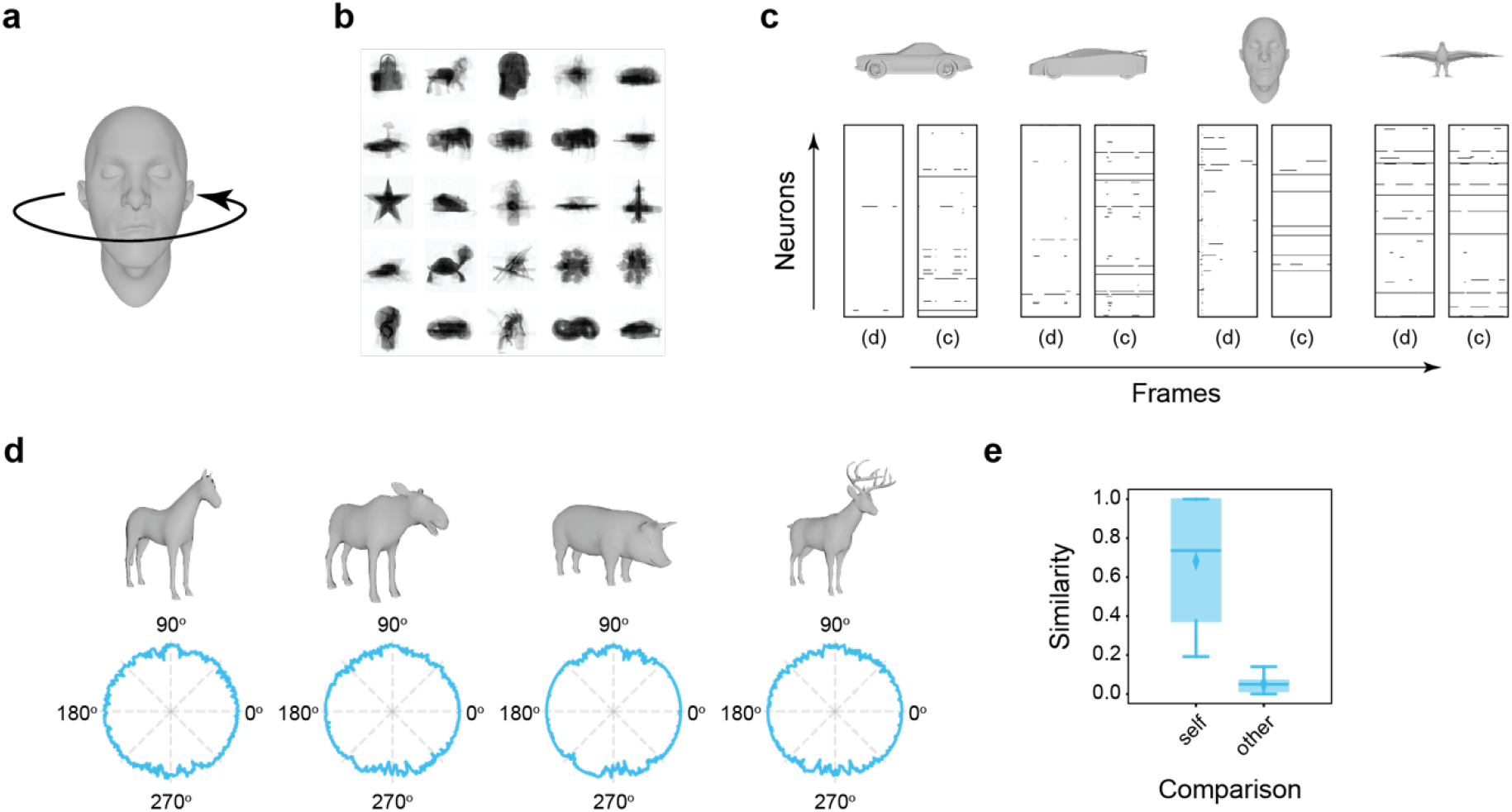
Perspective invariant representation of 3D objects. **a.** View of ration of 3D objects along the vertical axis is animated by take snapshots of individual angles sequentially. The network is exposed to short clips of these animations as in Fig. 4a. **b.** The network extracts complete and superimposed views of the objects in the discrimination layer. **c.** The responses of the discrimination (d) and classification (c) layer neurons to all views of the objects. Four examples are shown. Responses to the same object are consistent across different views in the classification layer. Note the distinct responses to two cars models. **d.** Representation consistency plots for 4 objects sharing similar characteristics. The representations are consistent for the same object. **e.** The average representation similarities between the frames originating from an object (self) were significantly higher than the representation similarities between frames of different objects (other).

### Manifold geometry of objects representation

Neuronal population response in the brain resides in a multi-dimensional space spanned by the activities of individual neurons^62–64^. Analytical methods can project these high-dimensional responses to low-dimensional manifolds^65–67^, along which we can measure perceptual distances for discrimination and classification purposes^68–71^. Recent theories propose that the manifolds’ geometry becomes more separable along the multiple sensory processing stages and gets straightened at later steps to allow invariant representations^10,72,73^. Indeed, psychoanalytical experiments indicate such transformations in the perceptual space^73^. In contrast, though ANNs show improved segregation between representations’ projections in their final layers, they fail to recapitulate the projection straightening observed early in the sensory processing^73,74^.

We analyzed the manifold structure of population response in our network. For rotating 3D objects, the low-dimensional manifolds in the pixel domain were jagged and occupied convoluted subspaces (Fig. 6a). The geometry becomes more organized in the discrimination module, with some examples occupying curved or rugged spaces (Fig. 6b). Strikingly, nearly all samples fall onto straightened hyperplane in the classification module, consistent with their invariant representation by the neurons. We assessed the linearity of representation manifolds in terms of their curvatures (see methods). With lower curvature indicating manifold straightening, considerable linearization was observed for all forms of variations in objects (Fig. 6c). Therefore, the transformation performed by our network has straightened the manifolds to allow perceptual invariance and robustness.

**Fig. 6.**
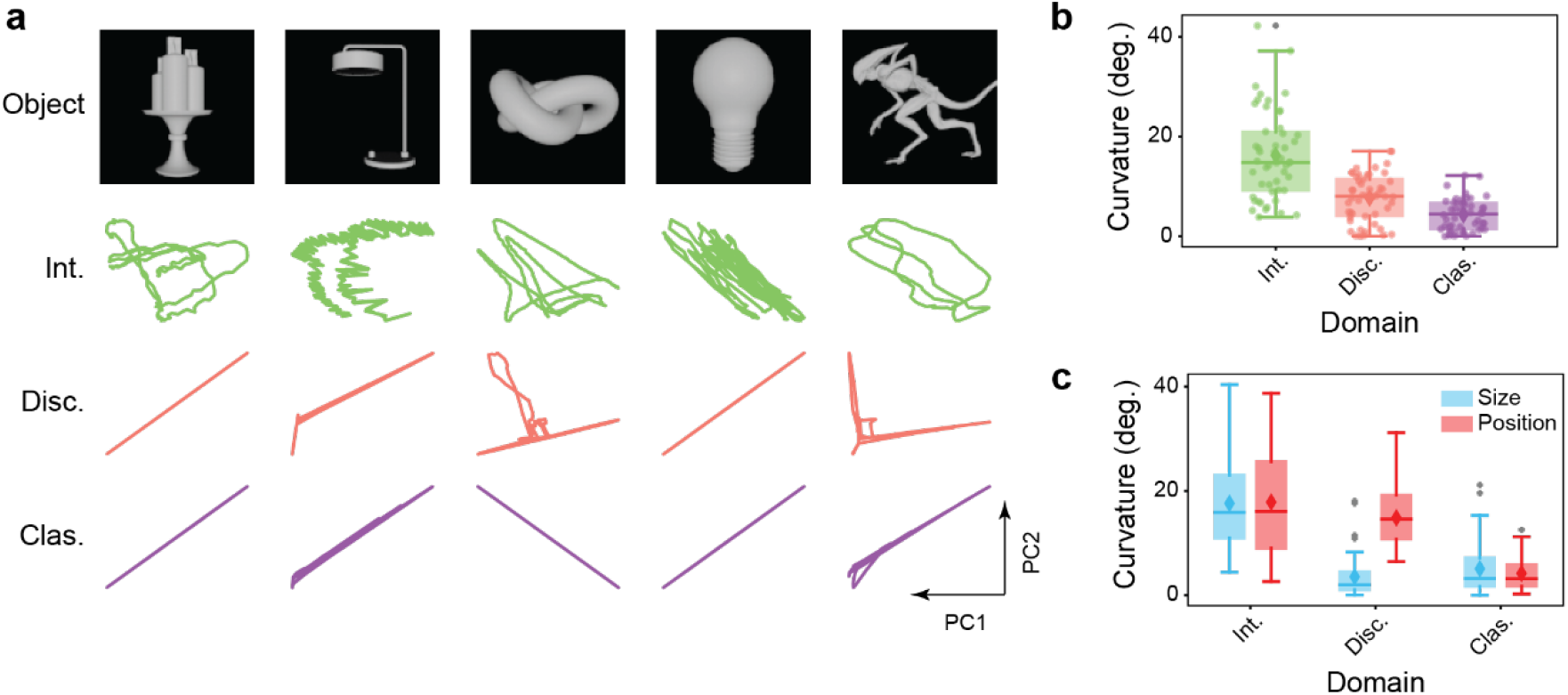
Manifold straightening in the network. **a.** Examples of 3D object models, their manifold structures in the intensity domain (Int), and the manifold structures of the representation in the discrimination (Disc) and classification (Clas) modules. Only the first two PCs are shown. **b.** Comparison of the average curvatures of the manifolds for all rotating objects in all domains. The manifolds are highly curved in the intensity domain but are progressively straightened. Manifolds of representations in the classification modules are straight. **c.** The same trend follows for size and position variations.

## DISCUSSION

The identity of an object is embedded in the structural relationships among its features. These relationships, or dependencies, can be utilized to encode object identity. A network that maximally captures these dependencies can identify the presence of an object without the accurate details of the input patterns^49^. Here, the specific configurations of our network allow dependence capturing to permit invariant representations. This network architecture is distinct from the hierarchical assembly model, which was first proposed by Hubel and Wiesel^75,76^ to explain the increasing complexity of receptive field properties along the visual pathway and later formed the foundation of convolutional neural networks^3^. These models assume that neurons in the cognitive centers recapitulate precise object details. Arguably, accurate object image reconstruction is not necessary for robust representation, and this deeply rooted assumption may have created unwanted complexity in modeling object recognition. Our model does not calculate reconstruction errors to assess its learning performance. By capturing dependencies that define objects and their classes, it produces remarkably consistent representations of the same object across different conditions. The size, translation, and rotation invariance show that the network can naturally link features that define an object or its class together without ostensibly being designed to do so. It permits the non-linear transformation of the input signals into a representation geometry suitable for identification and discrimination.

Importantly, our model illustrates how dependence capturing may allow the brain to learn about objects through local and continuous changes at individual synapses and stably represent them. The two circuit architectures are based on known connectivity patterns. Although both designs capture feature dependencies defining objects and classes, their connections differ and serve different functions. The discrimination module makes individual representations as distinctive as possible. The classification module binds class-specific features to highlight and distinguish different object types. This two-prong representation may give rise to perceptual distances that are not linearly related to the distances in input space.

Our model has profound implications for ANNs. It utilizes fundamentally different learning algorithms from the current models and does not rely on error propagation. It also avoids the problem of credit assignments in deep learning. As we have shown, the model can produce remarkable results that rival much more complicated networks with fewer neurons, fewer parameters, and no requirement for deep layers. Although these performances may be trumped by the highly sophisticated deep learning models that rely on superior computing power, our models can also be developed into complex structures to perform additional tasks with improved performance. Given that it requires much fewer examples to learn and is much more energy efficient, we expect ANNs based on the dependence capturing architecture to rival or outperform current ones.

## METHODS

### Generation of initial decorrelating connectivity

The current network is developed based on the principles of Maximal Dependence Capturing (MDC)^49^, which prescribes that individual neurons should capture maximum information about distinct objects. However, to achieve this, the network must be competent in differentiating objects in its initial response. In other words, the connectivity between the inputs and the representation neurons should allow distinct inputs to elicit disparate responses without specific learning. Random initial connectivity, which is commonly utilized in the models of object detection and classification, is not statistically inclined to induce such a response. We, therefore, introduced an initial bias in the connectivity that set the system to minimize the chances of co-activating two representation neurons. This ensured maximum distinction in the system’s initial response to various inputs. From an evolutionary perspective, this is akin to a neural circuit developed to accommodate the statistics of the input patterns in the environment.

Stated formally, we demanded the expected value of the variance-covariance matrix of the response profiles of neurons to be an identity matrix i.e.

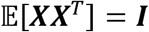

where ***X*** is the matrix of representations of different inputs and ***I*** is an identity matrix. Ignoring the non-linearity conferred to the system, we can approximate ***X*** in terms of input matrix ***Y*** and weight matrix ***ϕ*** as

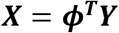

This relation gives

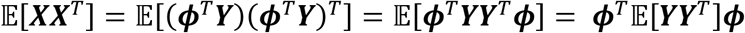

Clearly, 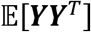 is the variance-covariance matrix of response profiles of input neurons (denoted by *∑_YY_*). With this relation, the above requirement of matching the variance-covariance matrix of representation neurons to the identity matrix reduces to solving the following equation

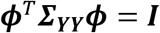

Since the variance-covariance matrix of any set of random variables is symmetric, it can be diagonalized using the orthogonal matrix ***Q*** of its eigenvectors i.e.

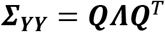

where ***Q*** is the matrix of orthogonal eigenvectors of ***∑_YY_*** and ***Λ*** is a diagonal matrix of eigenvalues of ***∑_YY_***. Using this transformation to solve the problem at hand, we have

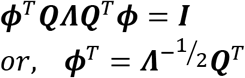

In general, any matrix 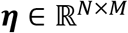 with orthogonal columns can be multiplied with the above solution, i.e.

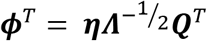

Thus, a connectivity matrix ***W***, as derived above, will make the variance-covariance matrix of representation neurons’ response profiles match the identity matrix.

### Learning rules in the discrimination module

Following the MDC principle, the main goal of the network is to allow the neurons to capture most informative structures from the inputs. Because these structures are not known *a priori*, the network should capture as much input structure as possible in distinct neuronal ensembles. To achieve this goal, the network updates its connections from the input layer to optimize the following function

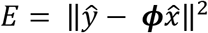

where *ŷ* is an input vector, 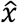 is its representational vector in the discrimination layer, and ***ϕ*** is the connectivity between the input and the discrimination layer. The updates in connectivity that optimize this function can be stated as

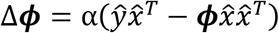

where *α* is the learning rate. Notably, the change in ***ϕ*** has two components, an additive component given by the rank one matrix 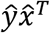, and a subtractive component given by the rank one matrix 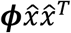. The additive part corresponds to the Hebbian update rule that strengthens the connection when an input neuron and a representation neuron fire together. Further, the (*i*, *j*) elementt of the matrix 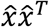 can be positive only when *x_i_* and *x_j_* are both positive. Thus, the second term of the update rule is anti-Hebbian in nature, i.e., the update reduces all the connections between input neurons and co-active representation neurons.

Note that the initial ***ϕ*** in the network is the generalized biased connectivity that ensures unique inputs induce distinct responses without any explicit constraints. It does not reflect input structures, and therefore, the matrix 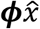 cannot be treated as the reconstruction of the input. The optimization procedure and the update rules only establish that comprehensive input features are learned on top of this decorrelating connectivity.

The recurrent weights *w* in the inhibition layer were set as

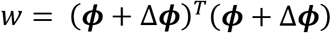

There was no normalization of ***ϕ*** before calculating recurrent weights. Therefore, if two neurons received similar input connections (similar columns of ***ϕ***), the symmetric inhibition between them was large. This increased the sparsity of representations and made them more unique.

### Update rules in the classification module

Weight between discrimination and classification neurons are updated based on the activities of the two neurons. The recurrent excitatory connections in the classification modules are initially set at 0, and all of neurons receive a global inhibition. The weights are updated based on the sum of potentiation between any pair of neurons. When two neurons get co-active together, their potentiation of connection increases. However, if only one of the two neurons gets active, then the potentiation of connection decreases. If both neurons remain inactive for a certain input, then their potentiation is unchanged i.e.

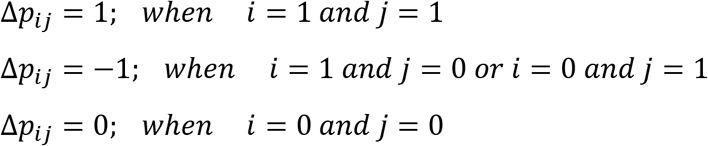

Here, Δ*p_ij_* is the change in the connection potentiation between neurons *i* and *j*. The connection weight is set as 1 if the sum of all potentials after encountering an arbitrary number of inputs crosses a set threshold. All other weights remain 0. The potentiation values of all possible connections are reset to zero, and the process of updating them restarts i.e.

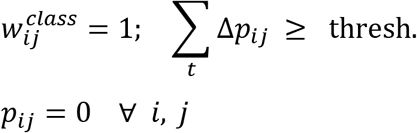

### Specificity calculations

To assess how specific an input’s representation is, we calculated the representation specificity. To estimate specificity, we first calculated the pairwise similarity between all representations of all objects and obtained a similarity matrix *S*. We then z-scored the similarity of an input’s representation to all other representations. In other words, we obtained the matrix *S_z_* as

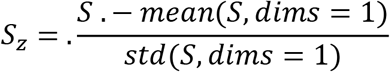

Here *mean*(*S, dims* = 1) an d *std* (*S, dims* = 1) denote the mean and standard deviation in the rows of the matrix *S*, and the dot operation (.) denotes elementwise calculations. The specificity of an input’s representation was its z-scored similarity with itself i.e.

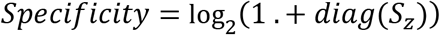

### Power spectrum calculations

To estimate the level of noise in images and their features learned by the network, we performed a power spectrum analysis on them. Both images and learned images were Fourier-transformed, and their log-power was calculated. The 2D log-power of the images and the learned structures were radially averaged to obtain the 1D power spectrum. The presence of noise was indicated by a higher power in higher frequencies of the spectrum. The comparisons were made using the highest 20% of the frequencies.

### Response consistency calculations

The representation of different views of 3D objects in the classification module consisted of neurons that are consistently active for all views of the object. We calculated the overall consistency of object representation in the classification module of the network. To calculate the consistency, we measured the cosine similarity between the representations of consecutive views of the object. The variation in the similarity indicated the consistency in representations. A lower variation in the similarity measures implied higher consistency and vice versa (see Extended Data Fig. 3 for details).

### Manifold generations and curvature calculations

To assess the geometry of manifold structures, we collected all views of all objects in the matrix I. Similarly, we collected their representations from discrimination and classification modules in matrices *R_d_* and *R_c_* respectively. We then performed Principal Component Analysis on all three matrices separately and plotted all views of individual objects as projections on the first two principal components. The plot depicted a 2D projection of the object manifolds.

To calculate the curvature of the 2D projection of the manifold, we selected three consecutive points *p_i_*, *p*_*i*+1_ and *p*_*i*+2_. We then measured the angle between vectors *p*_*i*+2_ – *p*_*i*+1_ and *p*_*i*+1_ – *p_i_* as

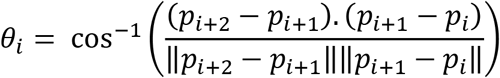

These angles were measured for all possible values of *i*. The curvature of the manifold was calculated as the average of all angle measures.

## Network sizes in figures

Fig. 1: The face images are 100 × 100 pixels and there are 500 discrimination layer neurons.

Fig. 2: a-c. Symbols are 16 × 16 pixels and discrimination layer size is 500. d-e. Both symbols and faces are 100 × 100 pixels and discrimination layer size is 1000.

Fig. 3: MNIST digits are 28 × 28 pixels, the discrimination and classification layers are both 10000-dimension.

Fig. 4–5; Extended Data Fig. 3: Object views are 100 × 100 pixels, representation sizes (both classification and discrimination) are both 1000.

Extended Data Figure 2: Same as Fig. 2d-e.

## Image and 3D object model sources

The symbols and facial images published previous^49^. 3D object models were downloaded from https://www.turbosquid.com.

## Author Contributions

CRY and RR developed key concepts and co-wrote the paper. RR performed analyses and modeling.

## Acknowledgments

This work fulfills, in part, requirements for RR’s Ph.D. thesis with the Open University, United Kingdom. We thank all Yu lab members for their insightful discussions and feedback.

## Extended Data

**Fig. 1.**
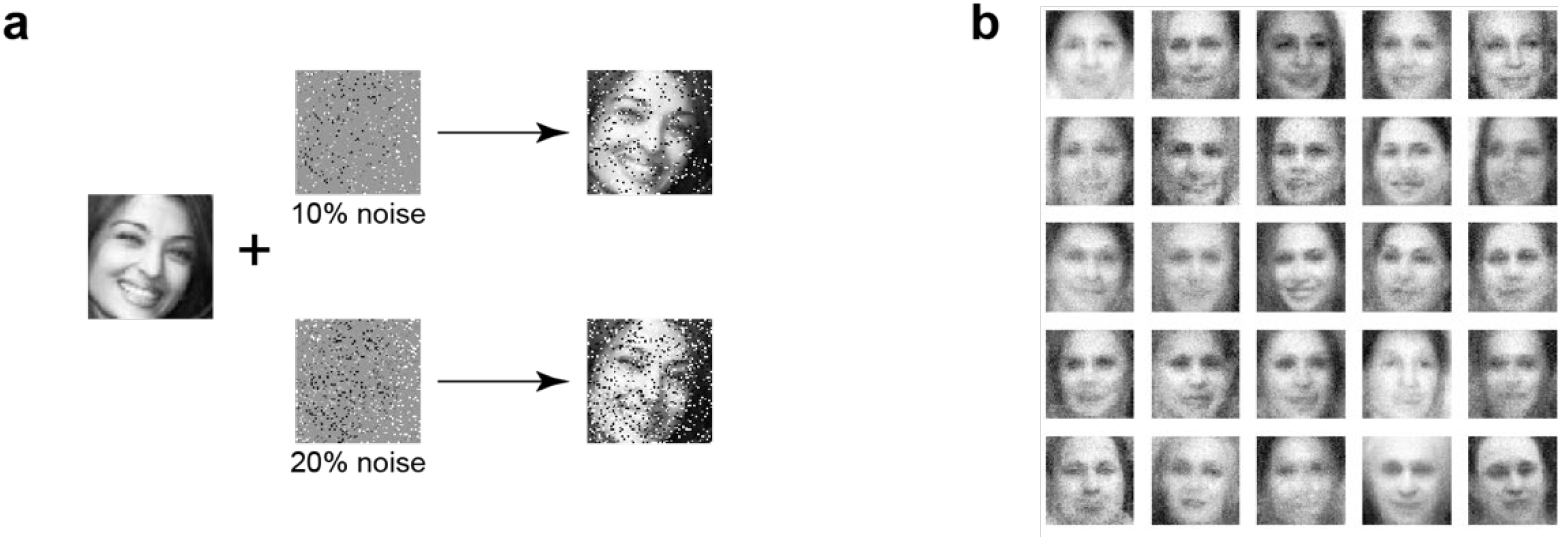
Generation of receptive fields. **a.** A face image with10% or 20% of pixels corrupted with a salt-and-pepper noise pattern. **b.** The receptive fields developed by the network from learning 20% noisy images. Images are from ref ^49^.

**Fig. 2.**
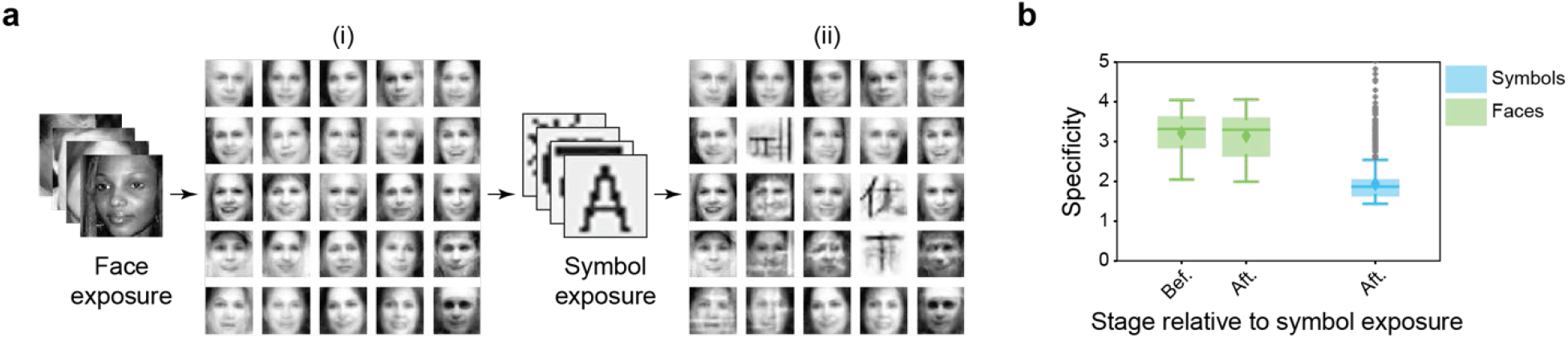
Continuous leaning. **a.** The network was exposed to symbols following limited learning of the faces. (i) The RFs after face exposure. (ii). Some RFs resembling symbols develop after the network is exposed to symbols. **b.** The specificity of face representations before and after symbol exposure is comparable. The representation specificity of symbols is also comparable to faces. Images are from ref ^49^.

**Fig. 3.**
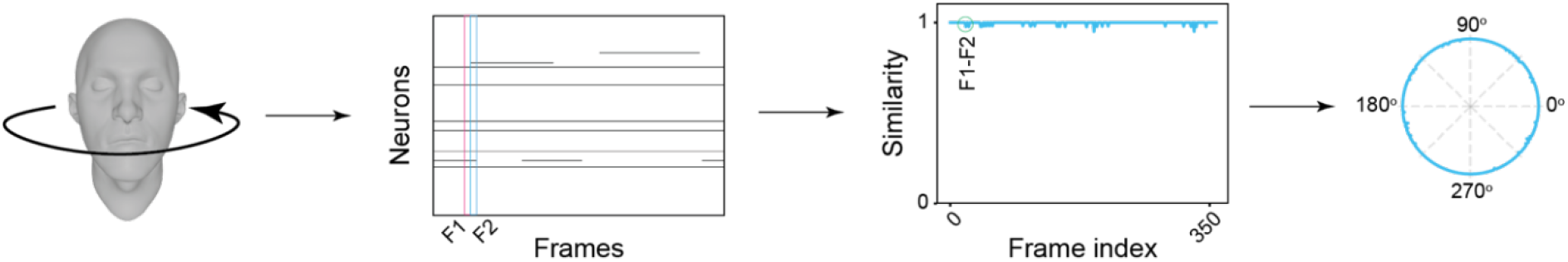
The process of generating representation consistency plots. A 3D object was rotated around the vertical axis and representations in the classification layer were recorded and displayed sequentially. Pairwise similarity between responses of two consecutive frames (illustrated as frames F1 and F2, magenta and cyan, respectively) were measured. By moving the frames down the sequence, similarity values were calculated account for all perspective angles. This similarity measured representational consistency. A value of 1 indicated identical responses between two frames. The circular plot, with a radius of 1, shows values corresponding to view angels.

